# Reversible inactivation of ferret auditory cortex impairs spatial and non-spatial hearing

**DOI:** 10.1101/2021.11.16.468798

**Authors:** Stephen M. Town, Katarina C. Poole, Katherine C. Wood, Jennifer K. Bizley

## Abstract

A key question in auditory neuroscience is to what extent are brain regions functionally specialized for processing specific sound features such as sound location and identity. In auditory cortex, correlations between neural activity and sounds support both the specialization of distinct cortical subfields, and encoding of multiple sound features within individual cortical areas. However, few studies have tested the contribution of auditory cortex to hearing in multiple contexts. Here we determined the role of ferret primary auditory cortex in both spatial and non-spatial hearing by reversibly inactivating the middle ectosylvian gyrus during behavior using cooling (n=2) or optogenetics (n=1). In optogenetic experiments, we utilized the mDLx promoter to express Channelrhodopsin 2 in GABAergic interneurons and confirmed both viral expression (n=2) and light-driven suppression of spiking activity in auditory cortex, recorded using Neuropixels under anesthesia (n=465 units from 2 additional untrained ferrets). Cortical inactivation impaired vowel discrimination in co-located noise, but not in clean conditions, or when the temporally coincident vowel and noise were spatially separated by 180°. Testing the ferrets implanted with cooling loops in a sound localization task confirmed that deficits in spatial hearing arose from inactivation of the same region of auditory cortex that was implicated in vowel discrimination in noise. Our results are consistent with single unit recordings in primary auditory cortex showing mixed selectivity for spatial and non-spatial features of sound and suggest a contribution of this region to multiple forms of hearing necessary for auditory scene analysis.

**Significance Statement:** Neurons in primary auditory cortex are often sensitive to the location and identity of sounds. Here we inactivated auditory cortex during spatial and non- spatial listening tasks using cooling, or optogenetics. Auditory cortical inactivation impaired multiple behaviors, demonstrating a role in both the analysis of sound location and identity and confirming a functional contribution of mixed selectivity observed in neural activity. Parallel optogenetic experiments in two additional untrained ferrets linked behavior to physiology by demonstrating that expression of Channelrhodopsin 2 permitted rapid light-driven suppression of auditory cortical activity recorded under anesthesia.

## Introduction

A central question in neuroscience is to what extent the brain is functionally organized into specialized units versus distributed networks of interacting regions (Földiák, 2009; Bowers, 2017). In sensory systems, separate cortical fields are thought to process distinct stimulus features such as visual motion, color and identity (Nassi and Callaway, 2009) or sound location and identity (Rauschecker and Scott, 2009).

Primary auditory cortex plays a critical role in many aspects of hearing. Neurons in this area show tuning to multiple features of sounds such as location and level (Brugge et al., 1996; Zhang et al., 2004), location and identity (Amaro et al., 2021) or vowel timbre, pitch and voicing (Bizley et al., 2009; Town et al., 2018). This sensitivity to multiple features can give rise to complex spectrotemporal tuning (Atencio et al., 2008; Harper et al., 2016) that can also be modulated by ongoing behavior (Fritz et al., 2003; David et al., 2012).

The mixed selectivity observed in responses of auditory cortical neurons is matched by a diverse range of behavioral deficits following auditory cortical lesions or inactivation (Slonina et al., 2022). Affected behaviors include discrimination of natural sounds such as vocalizations (Heffner and Heffner, 1986; Harrington et al., 2001), as well as sound modulation (Ohl et al., 1999; Ceballo et al., 2019) and sound localization (Heffner and Heffner, 1986; Malhotra et al., 2008). Most cortical inactivation studies focus on performance in a single task, or on a small range of related behaviors and thus our inferences on common functions must draw on data from different subjects, species and techniques.

Ideally, we would complement such inferences with direct comparisons of the effects of inactivation on distinct tasks performed by the same subjects and using the same methods of perturbation. Tests of distinct behaviors during auditory cortical inactivation are rare, but have yielded valuable insight into the functional specialization of non-primary auditory cortex (Adriani et al., 2003; Lomber and Malhotra, 2008; Ahveninen et al., 2013).

Here, we define distinct behaviors as those requiring subjects to act on the basis of orthogonal stimulus features, where orthogonality indicates that one feature can be varied while another remains constant (e.g. Flesch et al., 2018). The sparsity of inactivation data across distinct behaviors reflects the technical limitations on suppressing neural activity in humans and the difficulty in training individual animals to perform multiple tasks with contrasting demands.

We leveraged ferrets’ capacity to learn multiple psychoacoustic tasks to test the role of auditory cortex in distinct behaviors involving vowel discrimination in multiple contexts (clean conditions, or with co-located or spatially separated noise) and approach-to-target sound localization. During testing, we reversibly inactivated a large portion of primary auditory cortex by cooling the mid-to-low frequency area of the middle ectosylvian gyrus (MEG). Inactivation produced a pattern of deficits that confirms a common role for this brain region in both spatial and non-spatial hearing. Further experiments with optogenetics confirmed the role for MEG in vowel discrimination in noise and demonstrated the efficacy of light-driven suppression of sound evoked responses in ferret auditory cortex.

## Methods

### Animals

Subjects were ten pigmented ferrets (*Mustela putorius*, female, between 0.5 and 5 years old). Animals were maintained in groups of two or more ferrets in enriched housing conditions, with regular otoscopic examinations to ensure the cleanliness and health of ears.

Seven animals were trained in behavioral tasks in which access to water was regulated (**Table 1**). During water regulation, each ferret was water-restricted prior to testing and received a minimum of 60ml/kg of water per day, either during task performance or supplemented as a wet mash made from water and ground high-protein pellets. Subjects were tested in morning and afternoon sessions on each day for up to five days in a week, while their weight and water consumption was measured throughout the experiment.

**Table 1:** Metadata for each subject. Vowel discrimination was tested in clean conditions, with co-located (CL) noise, or spatially separated (SS) noise. Cooling loops implanted in F1216 (asterisk) were persistently blocked and could not be used reliably to achieve bilateral cooling (1 of 14 attempts). Animals implanted with microelectrodes provided single unit recordings for another study (Town et al., 2018).

| Ferret | Implant Type | Vowel Discrimination |  |  | Sound Localization |
| --- | --- | --- | --- | --- | --- |
|  |  | Clean | CL Noise | SS noise |  |
| F1311 | Cooling Loops | Yes | Yes | Yes | Yes |
| F1509 |  | Yes | Yes | Yes | Yes |
| F1201 | Microelectrode Arrays | No | No | Control only | No |
| F1203 |  | No | No | Control only | No |
| F1217 |  | No | No | Control only | No |
| F1216 | Cooling Loops* | No | No | Control only | No |
| F1706 | Optic Fibers | No | Yes | No | No |
| F1801 | Anesthetized Recording | No | No | No | No |
| F1807 |  | No | No | No | No |
| F1814 | Histology | No | No | No | No |

All experimental procedures were approved by local ethical review committees (Animal Welfare and Ethical Review Board) at University College London and The Royal Veterinary College, University of London and performed under license from the UK Home Office (Project License 70/7267) and in accordance with the Animals (Scientific Procedures) Act 1986.

### Stimuli

#### Vowel discrimination

Vowels were synthesized in MATLAB (MathWorks, USA) using an algorithm adapted from Malcolm Slaney’s Auditory Toolbox (https://engineering.purdue.edu/~malcolm/interval/1998-010/) that simulates vowels by passing a click train through a biquad filter with appropriate numerators such that formants are introduced in parallel. In the current study, four formants (F1-4) were modeled:/u/ (F1-4: 460, 1105, 2857, 4205 Hz), /ε/ (730, 2058, 2857, 4205 Hz), /a/ (936, 1551, 2975, 4263 Hz) and /i/ (437, 2761, 2975, 4263 Hz). Ferrets were only trained to discriminate between a pair of vowels: either /ε/ and /u/ (F1201, F1203, F1217, F1509 and F1706), or /a/ and /i/ (F1216 and F1311). All vowels were generated with a 200 Hz fundamental frequency.

Vowels were presented in clean conditions as two repeated tokens, each of 250 ms duration and of the same identity, separated with a silent interval of 250 ms (**Fig. 1A**). Here, two vowel tokens were used for consistency with previous work (Bizley et al., 2013a; Town et al., 2015). Sounds were presented through loudspeakers (Visaton FRS 8) positioned on the left and right sides of the head at equal distance and approximate head height. These speakers produced a smooth response (±2 dB) from 200 to 20000 Hz, with a 20 dB drop-off from 200 to 20 Hz when measured in an anechoic environment using a microphone positioned at a height and distance equivalent to that of the ferrets in the testing chamber. All vowel sounds were passed through an inverse filter generated from calibration of speakers to Golay codes (Zhou et al., 1992). Clean conditions were defined as the background sound level measured within the sound-attenuating chamber in which the task was performed in the absence of stimulus presentation (22 dB SPL).

**Figure 1.**
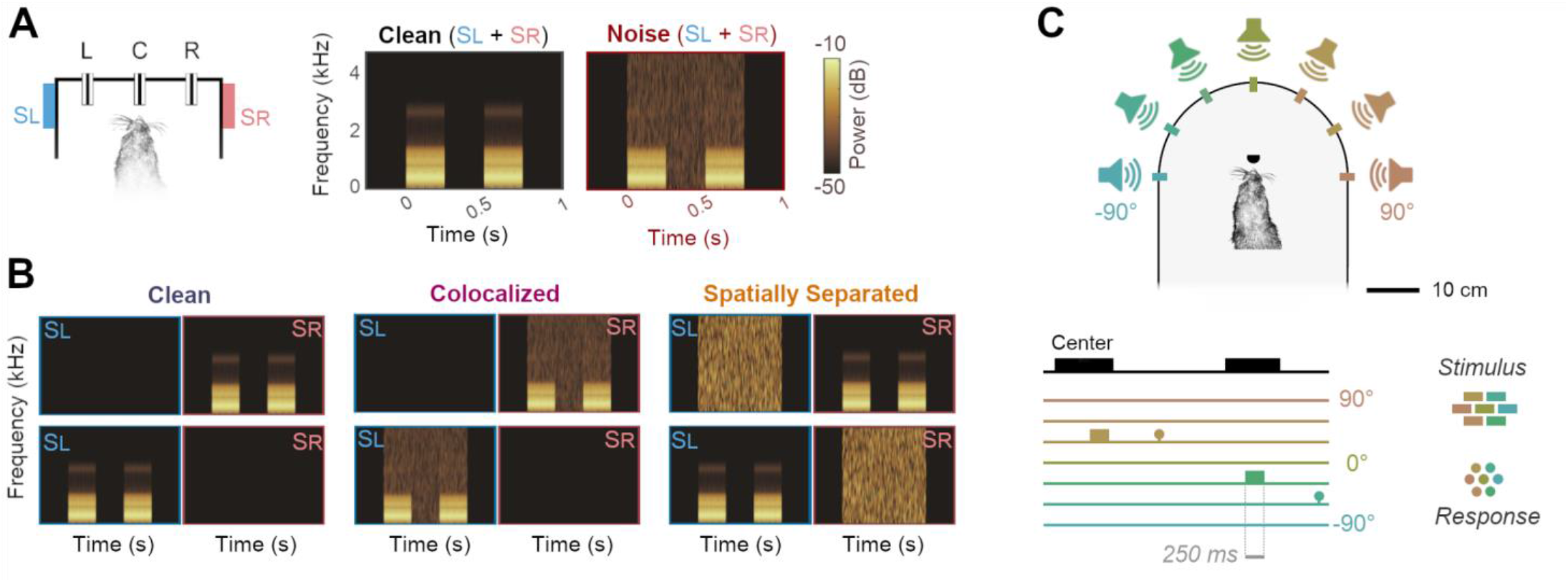
Behavioral task designs. **(A)** Vowel discrimination in noise and in clean conditions. Both vowel and noise were presented from speakers to the left (S_L_) and right (S_R_) of the head as the animal held at a center lick port (C). The animal then reported vowel identity by visiting either left (L) or right (R) response ports. Spectrograms show vowels (e.g. two 250 ms tokens of /u/, separated by 250 ms interval) alone or with additive broadband noise. Vowel identity was always the same for both tokens, and the animal was required to respond left or right based on that identity (i.e. there was no requirement to compare the two tokens). **(B)** Vowel discrimination task when vowels were presented from a single speaker in clean conditions, or with noise from the same speaker (colocalized) or the alternative speaker (spatially separated). Spectrograms and behavioral task arena as shown in A. **(C)** Sound localization task in which ferrets reported the location of broadband noise from one of several speakers in frontal space by approaching a water spout located at each speaker.

Vowels were also presented with additive broadband noise fixed at 70 dB SPL and restricted in time to the period of stimulus presentation (0 to 750 ms after onset of the first vowel token). Noise was generated afresh on each trial. Onsets of both vowels and noise were ramped using a 5 ms cosine function. During initial experiments on vowel discrimination in noise, vowels and noise were played from both left and right speakers (**Fig. 1A**); however when investigating spatial release from energetic masking, vowels were presented from either left or right speaker but not both. Noise was also presented from one speaker and thus the noise levels in such experiments were 67 dB SPL; noise was presented either from the same speaker as vowels, the opposite speaker or not at all (**Fig. 1B**).

#### Sound localization

Auditory stimuli were broadband noise bursts of differing durations (F1509: 700 ms; F1311: 250 ms or 100 ms) cosine ramped with 5-ms duration at the onset and offset and low-pass filtered below 22 kHz (finite impulse response filter <22 kHz, 70 dB attenuation at 22.2 kHz). Noise bursts were generated afresh on each trial in MATLAB at a sampling frequency of 48828.125 Hz and presented from one of seven speakers (Visaton FRS SC 5.9) positioned at 30° intervals (**Fig. 1C**). Note that one ferret (F1509) was not tested with sounds from the central speaker (0°). Across trials, stimuli were presented at one of three pseudo-randomly selected intensities (57, 61.5 and 66 dB SPL).

Speakers were calibrated to produce a flat response from 200 Hz to 25 kHz using Golay codes, presented in an anechoic environment, to construct inverse filters (Zhou et al., 1992). All the speakers were matched for level using a microphone positioned upright at the level of the ferret’s head in the center of the semi-circle. Calibrations were performed with a condenser microphone (Model 4191, Brüel and Kjær) and/or a Brüel and Kjær 3110–003 measuring amplifier.

### Task design

Behavioral tasks, data acquisition, and stimulus generation were all automated using custom software running on personal computers, which communicated with TDT real-time signal processors (Vowel discrimination: RZ6, Sound localization: RX8).

#### Vowel discrimination

Ferrets were trained to discriminate the synthetic vowel sounds within a custom-built double-walled sound attenuating chamber (IAC Acoustics Ltd.) lined with acoustic foam. The chamber contained a wire-frame pet-cage with three response ports housing infra-red sensors that detected the ferret’s presence. On each trial, the ferret was required to approach the center port and hold head position for a variable period (between 0 and 500 ms) before stimulus presentation. Animals were required to maintain contact with the center port until 250 ms after the presentation of the first token, at which point they could respond at left or right response ports. Correct responses were rewarded with water while incorrect responses led to a brief time-out (between 3 and 8 s) indicated by presentation of a 100 ms broadband noise burst and in which the center port was disabled so that animals could not initiate a new trial. Following a time-out, the animal was presented with a correction trial in which the same stimulus and trial parameters (e.g. hold time) were used. To suppress any bias the animal might have to respond at a particular port, we continued to present timeouts and correction trials until a correct response was made. Once a correct response was made on correction trials, a new vowel sound and trial parameters were selected for the next trial. To encourage animals to maintain a steady head position at the center port during sound presentation, a water reward was also given at trial onset on a small proportion (10%) of randomly chosen trials.

#### Sound localization

Ferrets were trained and tested in a second behavioral chamber that consisted of a custom-built D-shaped box surrounded by an array of seven speakers at 30° intervals. Each speaker had a response port located in front (8.5 cm in front of the speaker; 15.5 cm from the center of the box) at which animals could report sound location and obtain water rewards. A further port was also placed at the center of the arena to initiate stimulus presentation. This port was offset from the center by 3 cm to ensure the animal’s head was aligned at the center of the speaker ring, with the interaural axis in line with the −90° and +90° speakers. The distance between the head and speakers during sound presentation was 24 cm. Outside the training box, an LED (15 cm from the floor) was used to indicate trial availability. The test arena was housed in a custom built sound attenuating chamber (90 cm high x 90 cm wide x 75 cm deep, Zephyr Products Ltd, UK) lined with 45 mm acoustic foam.

### Behavioral training

#### Vowel discrimination

Subjects were trained to discriminate a pair of vowels through a series of stages of increasing difficulty. When first introduced to the training apparatus, animals were rewarded with water if they visited any port. Once subjects had associated the ports with water, a contingency was introduced in which the subject was required to hold the head at the central port for a short time (501–1001 ms) before receiving a reward. The central port activation initiated a trial period in which a nose-poke at either peripheral port was rewarded.

Following acquisition of the basic task structure (typically two to three sessions), sounds were introduced. On each trial, two repeats of a single vowel sound (each 250 ms in duration with a 250 ms interval) were played after the animal first contacted the port with a variable delay (between 0 and 500 ms). A trial was initiated if the subject’s head remained at the port for the required hold time, plus an additional 500 ms in which the first token of the sound and subsequent interval were played. Following trial initiation, vowel sounds were looped (i.e. played repeatedly) until the ferret completed the trial by visiting the “correct” peripheral port to receive a reward. Nose-pokes at the “incorrect” peripheral port were not rewarded or punished at this stage and incorrect responses did not terminate trials. If the animal failed to visit the correct port within a specified period after initiating a trial (between 25 and 60 s), that trial was aborted and the animal could begin the next trial.

Once animals were completing trials frequently, the consequences of incorrect responses were altered so that incorrect responses terminated the current trial. Subjects were then required to return to the center port to initiate a correction trial in which the same stimulus was presented. Correction trials were included to prevent animals from biasing their responses to only one port and were repeated until the animal made a correct response. After a minimum of two sessions in which errors terminated trials, a time-out (between 5 and 15 s) punishment was added to incorrect responses. Time-outs were signaled by a burst of broadband noise (100 ms), and the center port was disabled for the duration of the time out, preventing initiation of further trials.

Once subjects could discriminate repeated sounds on consecutive sessions with a performance of 80%, looping of sounds was removed so that subjects were presented with only two vowel sounds during the initiation of the trial at the center port. When ferrets correctly identified 80% of vowels in two consecutive sessions, the animal was considered to be ready for testing in noise. Note that beyond experience through testing, ferrets did not receive specific training to discriminate vowels in noise.

#### Sound localization

In contrast to vowel discrimination, training in sound localization took place after animals were implanted with cooling loops, and following completion of all testing in vowel discrimination. Ferrets (F1311 and F1509) were first trained to hold at the port in the center of the localization arena to initiate presentation of a series of repeating 1000 ms noise bursts (500 ms interval) from one speaker. The animal was allowed to leave the central port after the first burst, after which the stimulus repeated until a correct response was made at the peripheral port nearest the presenting speaker. Responses at other ports had no effect at this stage, but premature departures from the center triggered a short (1 sec) timeout.

Once ferrets were accustomed to the task (identified by regularly returning to the start port after receiving water from target locations), error detection was introduced so that trials were terminated when animals reported at the wrong peripheral port. The ferret was then required to initiate a new trial, on which the same stimulus was presented (correction trial) until a correct response was made. Time-outs were then introduced for incorrect responses and were increased from 1 to between 5 and 7 seconds. During this training phase, we also increased the hold time required at the central port before stimulus presentation, initially up to 500 ms during training and then 1000 ms during testing.

Once ferrets reached ≥ 60% correct, the stimulus was reduced to a single noise burst and subsequently the stimulus duration was reduced. Ferrets were ready for testing at these durations once their performance stabilized (approximately 3 to 4 weeks); for one ferret (F1311) we could reduce sound duration to between 250 and 100 ms with stable performance, however time constraints on the lifetime of the cooling implant required that we use a longer duration (700 ms) for the second animal (F1509). In all cases, animals were required to hold head position at the central port for the full duration of the sound and thus could not make head movements during stimulus presentation.

### Cortical inactivation using cooling

#### Loop implantation

Cortical inactivation experiments were performed using an approach developed by Wood et al. (2017): Two ferrets (F1311 & F1509) were successfully implanted with cooling loops made from 23 gauge stainless steel tubing bent to form a loop shape approximately the size of primary auditory cortex. (A third ferret, F1216, was also implanted but the loops were persistently blocked and thus non-functional). At the base of the loop, a micro-thermocouple made from twisting together PFA insulated copper (30 AWG; 0.254 mm) and constantan wire (Omega Engineering Limited, UK), was soldered and secured with araldite. Thermocouple wires were soldered to a miniature thermocouple connector (RS components Ltd, UK) and secured with epoxy resin prior to implantation.

Loops were surgically implanted over the middle ectosylvian gyrus, specifically targeting the mid-to-low frequency regions of primary auditory cortex (A1 and Anterior Auditory Field, AAF) that border the non-primary fields of the posterior Ectosylvian gyrus (**Fig. 2A**). Loops targeted this region as it is known to contain neurons sensitive to both sound timbre and location (Bizley et al., 2009; Town et al., 2018). Consistent with previous studies (Wood et al., 2017), we did not map the boundaries of auditory cortical subfields prior to loop placement. Cortical mapping may damage brain tissue, potentially triggering compensatory mechanisms that might mask a role in task performance. Placement of cooling loops was therefore based on our extensive experience targeting this area for electrode placements using anatomical landmarks (Bizley et al., 2009, 2013b; Town et al., 2018).

**Figure 2.**
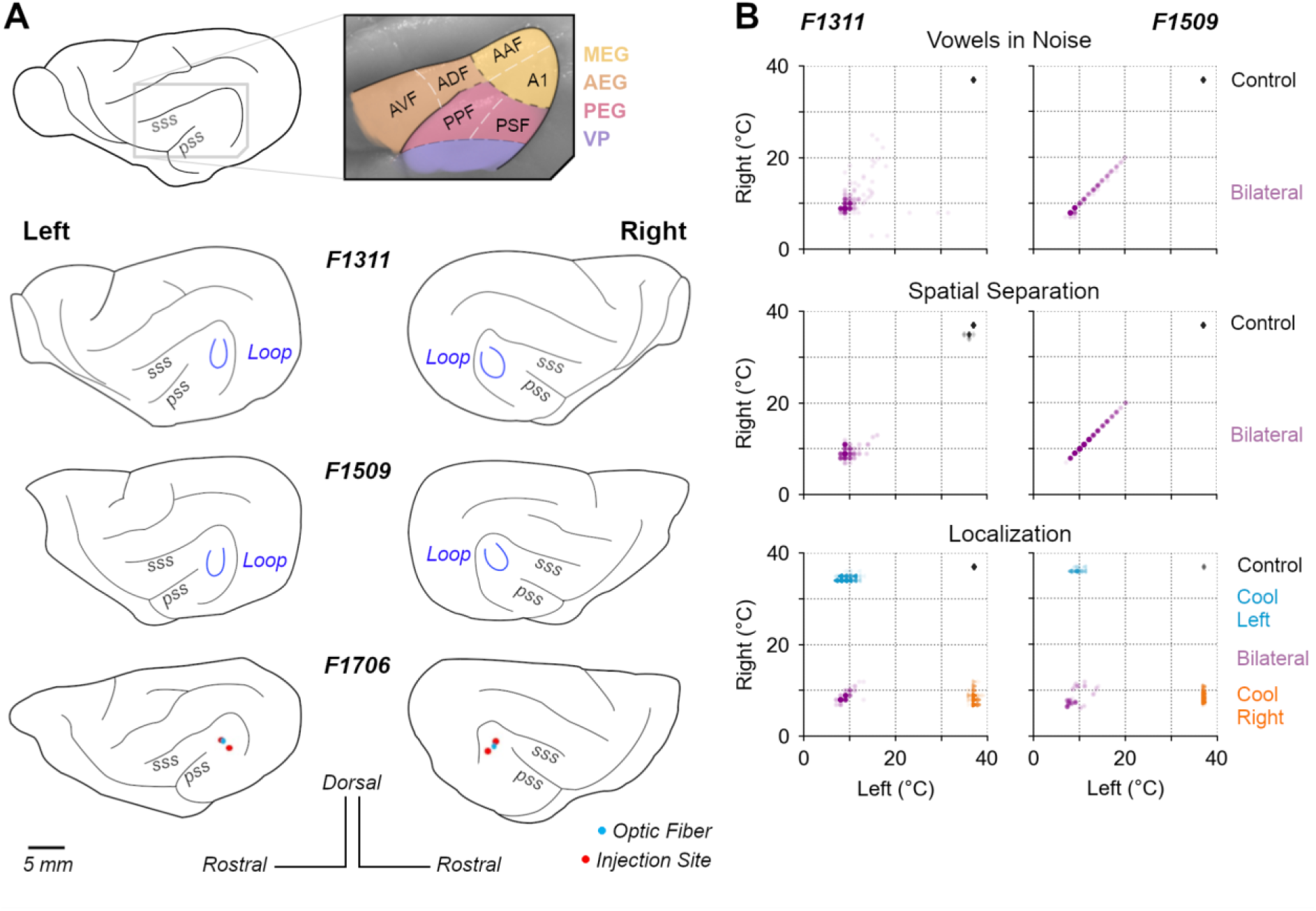
Cortical inactivation in behavioral tasks. **(A)** Anatomical location of ferret auditory cortex and positions of cooling loops (blue) implanted in F1311 & F1509 and viral injection in F1706 over the Middle Ectosylvian Gyrus. (Acronyms, A1: Primary auditory cortex, AAF: Anterior Auditory Field, AEG: Anterior Ectosylvian Gyrus, AVF: Anterior Ventral Field, ADF: Anterior Dorsal Field, PEG: Posterior Ectosylvian Gyrus, PPF: Posterior Pseudosylvian Field, PSF: Posterior Suprasylvian Field, VP: Ventral Posterior Auditory Field). **(B)** Distribution of cortical temperatures during bilateral cooling (all tasks) and unilateral cooling of left or right auditory cortex (sound localization only). Scatterplots show temperatures on individual trials measured at the base of each cooling loop, where contact was made with the cortical surface.

Surgery was performed in sterile conditions under general anesthesia, induced by a single intramuscular injection of diazepam (0.4 ml/kg, 5 mg/ml; Hameln) and ketamine (Ketaset; 0.25 ml/kg, 100 mg/ml; Fort Dodge Animal Health, Kent, UK). Animals were intubated and ventilated, and anesthesia was then maintained with between 1 and 3% isoflurane in oxygen throughout the surgery. Animals were provided with subcutaneous injections of atropine (0.09 ml/kg, 600 μl/ml) and dexamethasone (0.25 ml/kg), as well as surgical saline intravenously, while vital signs (body temperature, end-tidal CO_2_, Oxygen saturation and electrocardiogram) were monitored throughout surgery.

General anesthesia was supplemented with local analgesics (Marcaine, 2 mg/kg, AstraZeneca) injected at the point of midline incision. Under anesthesia, the temporal muscle overlying the skull was retracted and largely removed, and a craniotomy was made over the ectosylvian gyrus. The dura over the gyrus was then opened to allow placement of the cooling loop on the surface of the brain. The loop was shaped during surgery to best fit the curvature of the cortical surface prior to placement, and was then embedded within silicone elastomer (Kwik-Sil, World Precision Instruments) around the craniotomy, and dental cement (Palacos R+G, Heraeus) on the head. Bone screws (stainless steel, 19010-100, Interfocus) were also placed along the midline and rear of the skull (two per hemisphere) to anchor the implant. Implant anchorage was also facilitated by cleaning the skull with citric acid (0.1 g in 10 ml distilled water) and application of dental adhesive (Supra-Bond C&B, Sun Medical). Some skin was then removed in order to close the remaining muscle and skin smoothly around the edges of the implant.

Pre-operative, peri-operative and post-operative analgesia and anti-inflammatory drugs were provided to animals under veterinary advice. Animals were allowed to recover for at least one month before resuming behavioral testing and beginning cortical inactivation experiments.

#### Cooling during behavior

To reduce the temperature of the cortical tissue surrounding the loop, cooled ethanol (100%) was passed through the tube using an FMI QV drive pump (Fluid Metering, Inc., NY, USA) controlled by a variable speed controller (V300, Fluid Metering, Inc., NY, USA). Ethanol was carried to and from the loop on the animal’s head via FEP and PTFE tubing (Adtech Polymer Engineering Ltd, UK) insulated with silicon tubing and, where necessary, bridged using two-way connectors (Diba Fluid Intelligence, Cambridge, UK). Ethanol was cooled by passage through a 1 meter coil of PTFE tubing held within a Dewar flask (Nalgene, NY, USA) containing dry ice and ethanol. After passing through the loop to cool the brain, ethanol was returned to a reservoir that was open to atmospheric pressure.

For a cooling session, the apparatus was first ′pre-cooled’ before connecting an animal by pumping ethanol through spare cooling loops (i.e. loops that were not implanted in an animal) until loop temperatures fell below 0°C. The animal was then connected to the system, using the implanted thermocouples to monitor loop temperature at the cortical surface. The temperature was monitored online using a wireless transfer system (UWTC-1, Omega Engineering Ltd., UK) or wired thermometer, and pump flow rates adjusted to control loop temperature. Loops over both left and right auditory cortex were connected during bilateral cooling (all tasks), whereas only the left or right loop was connected during unilateral cooling (sound localization only).

For F1311, the animal was connected to the system and cooling began before the behavioral session, with the subject held by the experimenter and rewarded with animal treats (Nutriplus gel, Virbac, UK) while cooling pumps were turned on and loop temperatures reduced over five to ten minutes. When loop temperatures reached ≤ 12°C, the animal was placed in the behavioral arena and testing began. In contrast, F1509 would not perform tasks after being rewarded by the experimenter and so behavioral sessions were started and cortical temperature slowly reduced during task performance. Trials performed before the loops reached ≤ 20°C were excluded from analysis. Across animals, we considered bilateral cooling to be successful on any trial on which the average temperature across left and right loops was ≤ 20°C, while successful unilateral cooling required that one loop remained uncooled (≥ 35°C) while the other was ≤ 20°C (**Fig. 2B**). Cooling to these temperatures should suppress spiking activity within the immediate vicinity of the loop without spreading beyond the ectosylvian gyrus (Lomber et al., 1999; Coomber et al., 2011; Wood et al., 2017).

For both animals, cooling took place while the animals were free to move without interaction with the experimenter and within the same apparatus used for previous behavioral testing. The behavioral tasks during cooling were unchanged from those already described; i.e. the same ranges of sound levels were used, correction trials were included and the same reward contingencies were used. For each trial in the task, the time of stimulus onset was recorded and cross-referenced with temperature records so that any trials in which cortical temperature was above threshold during a cooling session could be removed from the analysis. During control testing, animals were connected to the cooling system using the same thermocouple sensors, but cooling loops were not connected to FEP tubing in order to avoid blockages and maximize the functional lifespan of loops.

### Data analysis: Behavior

All analyses excluded responses on correction trials, or trials where ferrets failed to respond within the required time (60 s). For all tests of vowel discrimination, we also required a minimum number of trials (n=10) and sessions (n=3) in both cooled and control conditions to include a sound level or SNR value in the analysis. Note that the requirement for a minimum number of trials introduced slight differences in the range of levels or SNRs tested between vowel discrimination experiments using vowel presentation both from left and right speakers and spatial release from energetic masking.

Temperature measurements were obtained on each trial for loops over left and right auditory cortex, and the animal was considered to be cooled if the average loop temperature was ≤ 20°C (bilateral cooling). In unilateral cooling, cooling was considered to be achieved if the relevant loop was ≤ 20°C. The threshold for cooling was based on previous work demonstrating the suppression of neural activity below this temperature (Jasper et al., 1970; Lomber et al., 1999).

Statistical analysis of effects of stimulus manipulation (e.g. presence of noise) and cooling used generalized linear mixed models (GLMMs) fitted using *lme4* (Bates et al., 2015) in R (version 4.2.1). The details of each model are outlined alongside the relevant results; however, in general, analysis of behavioral performance (correct vs. incorrect responses) was based on logistic regression in which the GLMM used binomial distribution and logit link function settings. For each model, we used ferret as a random factor and reported the magnitude of coefficients (β) of fixed effects of interest (e.g. effect of cooling) and probability (p) that the coefficient was drawn from a distribution centered about zero. To check model fit, we used the *DHARMa* package to assess the randomized quantile residuals (Dunn and Smyth, 1996) and reported both the marginal and conditional R^2^ values (Nakagawa and Schielzeth, 2013).

#### Vowel Discrimination

To analyze the effects of cooling, we compared behavioral performance of each animal across multiple sessions: The effects of cooling were measured on paired testing sessions performed on the same day (F1509) or unpaired sessions collected over the same time period (F1311). For F1509, we excluded trials when the animal was tested with sound levels below 50 dB SPL, for which no other subject was tested.

To summarize performance of each subject in a particular stimulus condition (clean conditions, co-located noise etc.), we randomly resampled (bootstrapped) data with equal numbers of each vowel and sound level or SNR (when showing data across level or SNR). Bootstrapping was performed 10^3^ times, with samples drawn with replacement on each iteration. For each bootstrap iteration, the number of samples drawn for each sound level or SNR was defined by taking the median of the number of trials sampled at each level or SNR. (For example, if we originally collected 10, 20 and 30 trials at 50, 60 and 70 db SPL, we randomly drew 20 trials with replacement for each sound level). Where vowels varied in sound location, we also resampled with equal numbers of trials with vowels from left and right speakers.

#### Sound Localization

Performance localizing sounds was analyzed using the percentage of trials on which animals correctly reported the target stimulus position. For F1311, we included responses to sounds of 100 ms and 250 ms duration, and sampled a random subset of data to ensure equal numbers of trials with each sound duration were included for each cooling condition. For each animal, we considered control data from all sessions after training was complete, and all trials obtained during cooling. When bootstrap resampling, we randomly drew equal numbers of trials when sounds were presented at each location (F1311: 69 trials at each of seven locations; F1509: 27 trials at each of six locations).

### Optogenetics

#### Injections in Auditory Cortex

Four ferrets (F1706, F1801, F1807 and F1814) were injected bilaterally in auditory cortex with an Adeno-associated Virus (AAV) to induce expression of Channelrhodopsin 2 (ChR2) in GABAergic interneurons using the mDlx promotor (AAV2.mDlx.ChR2-mCherry-Fishell3.WPRE.SV40, Addgene83898, UPenn Vector Core)(Dimidschstein et al., 2016). For each auditory cortex (i.e. left and right), injections were placed at two sites in the same area of MEG in which cooling loops were placed, under general anesthesia using the same sterile surgical protocol as described above. Within each site, injections were made at two depths (500 and 800 μm below the cortical surface) so that a total of four injections were made, with 1 μl injected each time.

#### Optogenetic testing during behavior (F1706)

Following viral delivery, we implanted an optrode (Neuronexus, Ann Arbor, MI, USA) in each auditory cortex to deliver light in F1706. During testing, light was delivered from a 463 nm DPSS laser (Shanghai Laser & Optics Century Ltd. China) with a steady-state power of 40 mW, measured at fiber termination before the optrode using an S140C integrating sphere photodiode sensor (ThorLabs, Germany). Although the optrode implanted included recording sites for monitoring neural activity during testing, we were unable to eliminate grounding issues that made recordings from this animal unusable and we therefore elected to train the animal in the vowel discrimination task and look for behavioral effects of silencing auditory cortex. The optrode was housed within an opaque plastic tower (25 mm tall) embedded in dental cement.

Retraining and testing of this animal after viral injection and optrode implantation was delayed due to the Covid-19 pandemic and behavioral testing took place 20 months after injection. At this point, we were only able to test the effect of light delivery on vowel discrimination in noise and a subsequent failure in the implant precluded testing of vowel discrimination in clean conditions, or with stimuli used to study spatial release from energetic masking or sound localization. The implant failure also prevented us from perfusing the brain of this animal in order to detect viral expression (although see below for successful confirmation of viral expression in other animals).

All data during vowel discrimination in noise was collected when the animal was attached to the optical fiber system, with opaque black tape used to secure the attachment and ensure that laser light was not visible to the ferret. In behavioral testing, light was delivered on 50% of test trials (with the exception of the first test session in which the laser was presented on all test trials); however, all correction trials took place without light delivery. On each trial that light was presented, we used short pulses (10 ms duration, presented at 10 Hz) that began 100 ms before sound onset, and continued until 100 ms after sound offset.

Data analysis for performance discriminating vowels in noise followed the same procedure as for analysis of behavior in animals with cooling. However, optogenetics provided more refined temporal control than cooling, allowing us to compare performance on trials within the same test session, with and without light delivery.

#### Optogenetic suppression of cortical activity (F1801 and F1807)

Photostimulation in visual cortex of ferrets expressing ChR2 in GABAergic interneurons suppresses cortical activity (Wilson et al., 2018). To determine if stimulation of ChR2 in GABAergic neurons was also sufficient to suppress sound-driven responses in auditory cortex, we recorded the activity of auditory cortical neurons while presenting stimuli with and without laser stimulation to ferrets under anesthesia.

Anesthesia was induced by a single dose of ketamine (Ketaset; 5 mg/kg/h; Fort Dodge Animal Health) and medetomidine (Domitor; 0.022 mg/ kg/h; Pfizer). The left radial vein was cannulated and anesthesia was maintained throughout the experiment by continuous infusion (ketamine: 5 mg/kg/hr; medetomidine: 0.022 mg/kg/hr; atropine sulfate: 0.06 mg/kg/hr and dexamethasone: 0.5 mg/kg/hr in Hartmann’s solution with 5% glucose). The ferret was intubated, placed on a ventilator (Harvard Model 683 small animal ventilator; Harvard Apparatus) and supplemented with oxygen. Body temperature (38°C), electrocardiogram and end-tidal CO_2_ were monitored throughout the experiment (^~^48 hours).

Animals were then placed in a stereotaxic frame and the site of viral injection over both left and right auditory cortex was exposed. A metal bar was attached to the midline of the skull, holding the head without further need of a stereotaxic frame. The animal was then transferred to a small table in a soundproof chamber (Industrial Acoustics, Winchester, UK) for stimulus presentation and neural recording. During recordings, the craniotomy was covered with 3% agar, replaced at regular intervals.

Neural activity was recorded in SpikeGLX (v 3.0., billkarsh.github.io/SpikeGLX) using Neuropixels Probes (IMEC, v1.0) inserted orthogonal to the cortical surface, and connected via headstages to an IMEC PXIe data acquisition module within PCI eXtensions for Instrumentation (PXI) hardware (PXIe-1071 chassis and PXI-6132 I/O module, National Instruments) that sampled neural signals at 30 kHz. Candidate action potentials were then extracted and sorted in Kilosort (v3.0., github.com/MouseLand/Kilosort), and manually curated to identify single (n = 174) or multi-unit (n = 291) activity. Spike clusters were merged based on assessment of waveform similarity and classed as a single unit using waveform size, consistency and inter-spike interval distribution (all single units had ≤2% of spikes within 2ms). Spike clusters without negative deflections in the waveform, which were primarily laser artifacts (identified as sharp peaks in the waveforms) were discarded from the analysis.

During recording, we presented broadband noise bursts of varying levels (40 to 70 dB SPL) and durations (50, 100 and 250 ms), either alone or with laser on. Stimuli were repeated 20 times, with a pseudo-random interval (0.5 to 0.7 seconds) between trials. Laser stimulation was provided by the same 463 nm DPSS laser used in behavioral experiments with F1706, attached to a custom made optic-fiber (1.5 mm diameter, Thorlabs FP1500URT) that was designed to maximize the area over which light was delivered, and could provide up to 300 mW at the fiber tip. Here, we report effects of pulsed light, delivered with a target power of 50 mW and frequency of 1 or 10 Hz. Pulses had a square-wave design with 50% duty cycle, beginning 100 ms before sound onset and ending 100 ms after sound offset. In addition to laser testing with sound presentation, we also tested the effect of the laser on spontaneous activity without sound. The effects of laser light delivery were measured at several sites over auditory cortex by placing the optic fiber and Neuropixel probe in various configurations over MEG and close to the viral injection sites of auditory cortex in each animal.Due to the COVID-19 pandemic, recordings were delayed until 21 months (F1801) and 18 months (F1807) after viral injection.

After recordings were completed, each animal was transcardially perfused with 0.9% saline and 4% paraformaldehyde (PFA) under anesthesia. The brain was then removed for storage in PFA, before sinking in 30% sucrose for 5 days prior to cryosectioning. Due to unavailability of a functioning cryostat (also delayed by the pandemic), brains were stored in PFA for six months, potentially limiting the quality of fluorescent signals. Coronal sections (50 μm) were taken through the full extent of the ectosylvian gyrus in order to confirm viral expression via visualization of mCherry. To better judge the quality of viral expression on a shorter timescale, we also transcardially perfused a further animal (F1814) within 12 weeks of viral injection, sectioned it immediately and measured mCherry and cell body (DAPI) labeling. Slices were imaged using a Zeiss Axio Imager 2.0 and Zeiss Confocal, and processed on Zen Blue.

### Data analysis: Optogenetic modulation of neural activity

To contrast the effects of laser light delivery on sound-driven activity, we first calculated the mean firing rate of each unit during auditory evoked activity, taking a window from sound onset to sound offset (50, 100 or 250 ms in length). For each unit, we compared the mean firing rate during this window calculated over all conditions in which the laser was present with the mean firing rate when the laser was absent (change in firing rate = laser OFF - laser ON). To contrast the effects of laser light delivery on spontaneous activity of each unit, we performed the same calculation on mean firing rates during the 100ms window before sound onset, on trials with and without the laser.

Inspection of neural activity with, and without laser suggested that light delivery had distinct effects on subgroups of neurons. To test if units could be distinguished by their modulation to laser delivery and to determine the number of separable groups of units using an unsupervised approach, we applied K-means clustering to the firing rates of each unit with and without laser. Clustering was based on the cosine distance between units (rather than Euclidean distance) in order to isolate the change in spike rate with laser stimulation across units with widely varying baseline firing rates. We identified the appropriate number of clusters within the data by comparing the sum of point-to-centroid distances for K = 1 to 10 and finding the knee-point using vector bisection (Dmitry Kaplan 2022. Knee Point, MATLAB Central File Exchange. https://www.mathworks.com/matlabcentral/fileexchange/35094-knee-point).

To map the extent of sound-evoked activity across the length of the probe, we compared mean spike rates during sound presentation and a time window preceding sound onset of matched duration (Wilcoxon signed-rank test). This analysis was performed on each unit to each sound duration by sound level condition, with Bonferroni correction for multiple comparisons. Units that showed a significant response in any of the conditions were classed as an auditory evoked unit (n = 72). We then contrasted the effects of laser light delivery on the firing rates of units recorded at different cortical depths during sound presentation, where depth refers to the distance on the probe from the most superficial channel on which spiking activity was observed.

We also investigated the temporal dynamics of the optogenetic stimulation to control for heating effects from laser delivery (Owen et al., 2019). To identify the latency at which light delivery induced a significant change in firing, we performed nonparametric cluster statistical analysis, which controls for multiple comparisons that would occur from calculating a test-statistic over each timepoint, by calculating a test-statistic from clusters of adjacent time samples of the PSTH in which firing rate with laser was greater than without laser (or vice versa)(Maris and Oostenveld, 2007). This statistic was calculated during the 100 ms after laser onset for each condition and the minimum time bin labeled as significant by the cluster statistic was averaged across conditions to calculate the latency for each unit.

## Results

### Optogenetic inactivation of sound-driven responses in auditory cortex

We used an AAV vector with an mDlx promoter to target expression of ChR2 to GABAergic interneurons in ferret auditory cortex. Post-mortem histology confirmed viral expression in two of three animals perfused (F1807 and F1814, but not F1801, in whom terminal recordings had severely compromised brain quality). Widefield imaging demonstrated viral expression in MEG, with labeled cells observed up to between 1 and 2 mm from injection sites (**Fig. 3A**). Confocal imaging revealed colocalization of mCherry with cell bodies (labeled by DAPI), with the opsin localized around the cell body (F1814).

**Figure 3.**
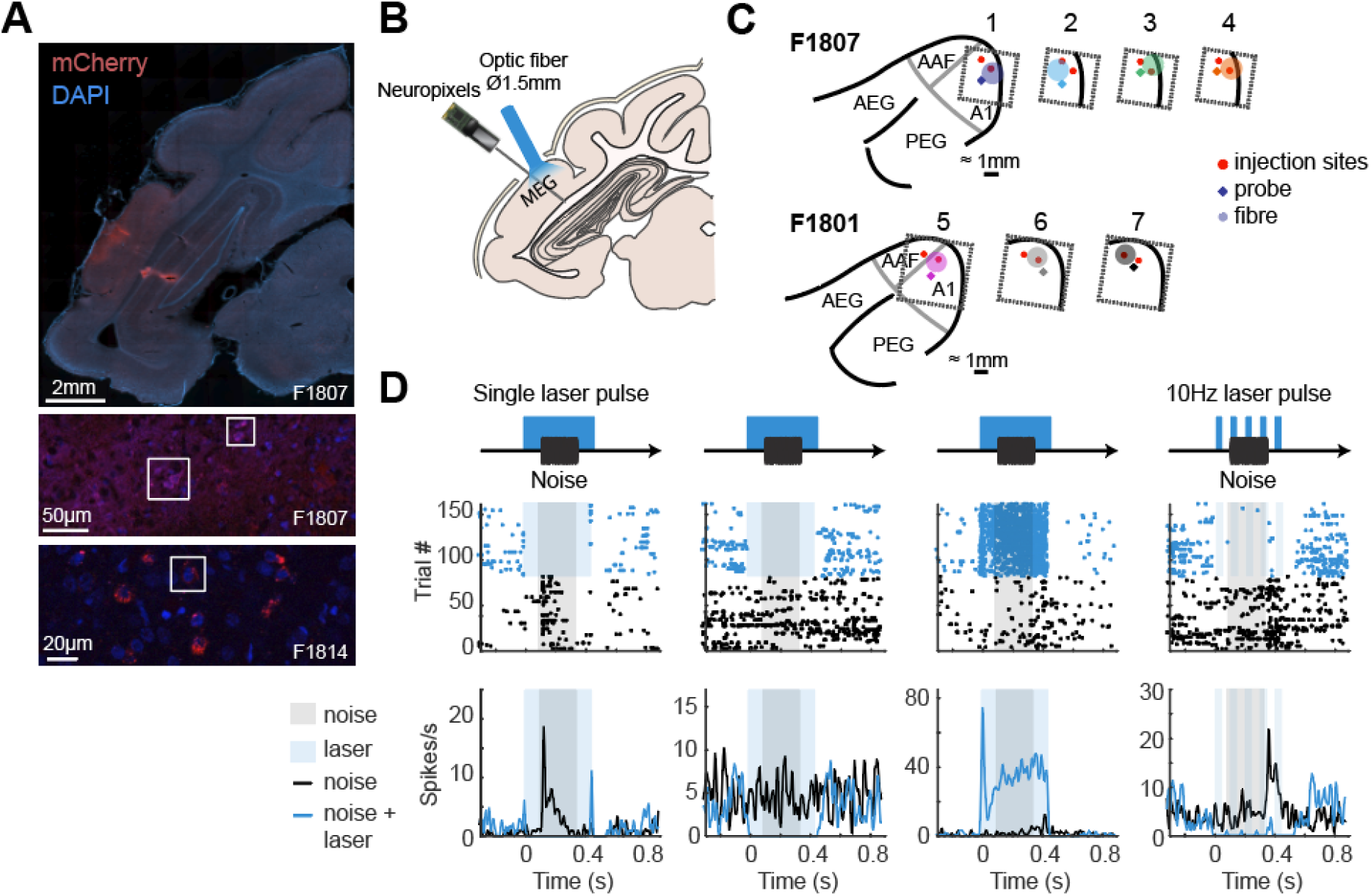
Targeting of optogenetic inactivation and neural responses. (**A**) Imaging viral expression in ferret auditory cortex. Top: Widefield imaging of coronal sections through the ectosylvian gyrus with the cell bodies labeled with DAPI (blue) and ChR2 labeled with mCherry (red). Middle / Bottom: Confocal imaging of the injection site showing colocalization of cell bodies and mCherry expression (outlined). (**B**) Experimental schematic showing stimulus and light delivery protocols. (**C**) Configurations of probe and optic fiber over injection sites within MEG in each ferret (F1807 and F1801). (**D**) Peri-stimulus time histogram and raster plots showing responses of four example units recorded from auditory cortex with and without light delivery in a single laser pulse (columns 1-3) and a 10 Hz laser pulse (column 4).

We then examined the electrophysiological efficacy of cortical inactivation through optogenetics using Neuropixels probes to record the activity of 465 units (n = 174 single units, 291 multi-units) in auditory cortex under ketamine-medetomidine anesthesia. Multiple optic fiber and recording sites were tested over auditory cortex, and at each site, we presented broadband noise with half of the trials having a laser delivery simultaneously presented (from 100 ms before, to 100 ms after sound onset/offset; **Fig. 3B-C**). Light delivery affected neural responses in a variety of ways, including suppressing responses to sound, suppressing baseline spontaneous firing and, in some cases, driving firing (**Fig. 3D**).

When comparing the mean firing rates of units with and without laser light, we saw that the activity of most units were suppressed by light, but that a small number showed no change in firing, or increases in firing. While these patterns were most evident when examining firing in the time window around sound presentation (**Fig. 4A**), the same pattern was also evident in spontaneous activity (**Fig. 4B**). During spontaneous activity the lower firing rates of units gave less scope to observe modulation and thus the effects of inactivation were weaker.

**Figure 4.**
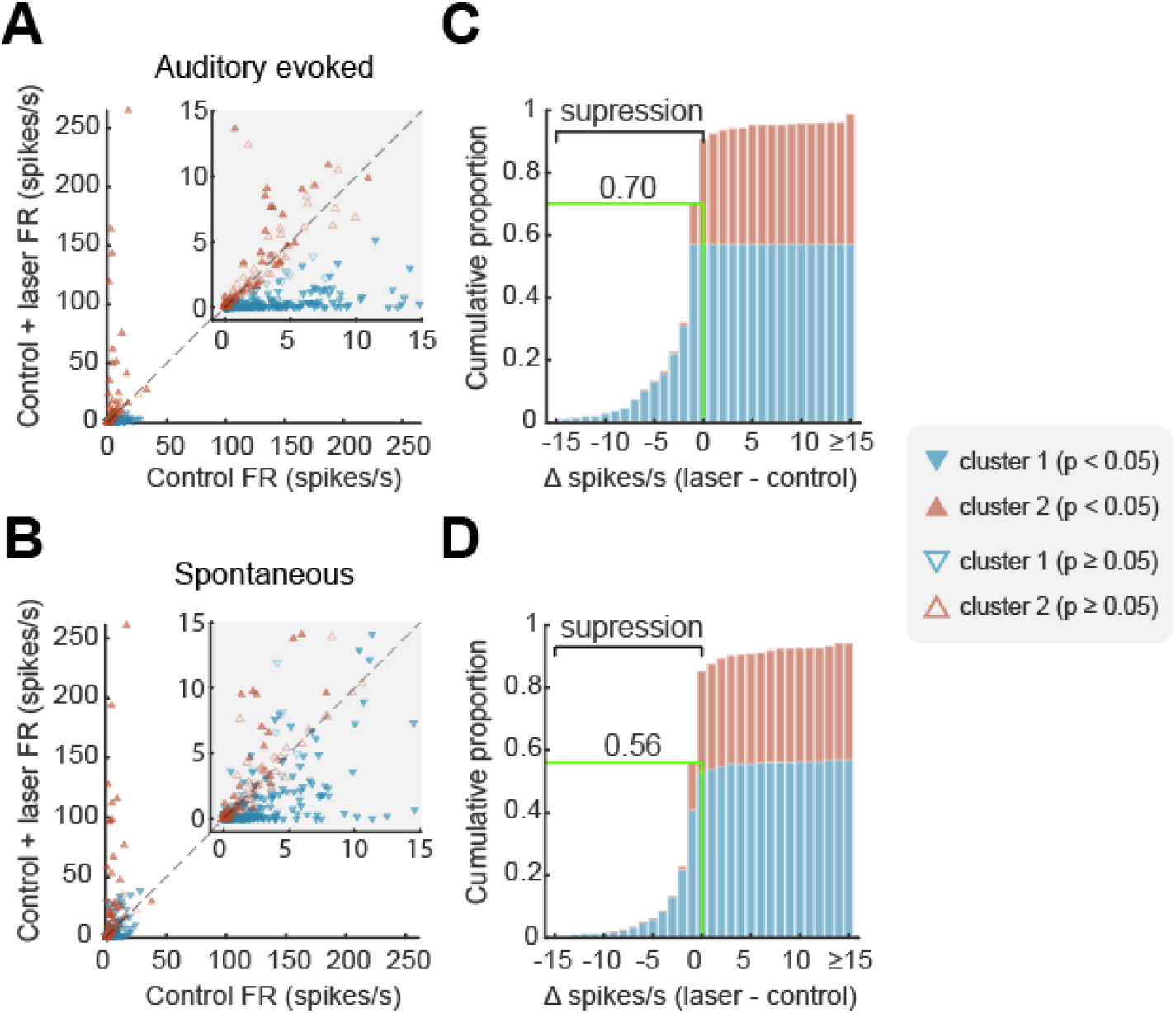
Optogenetic inactivation of auditory cortical activity. (**A-B**) Scatterplots of firing rate with and without laser and (**C-D**) cumulative histograms of change in firing rate with laser light delivery. Plots show firing rate measured during (**A, C**) or before (**B, D**) sound presentation for each unit, coloured by cluster and filled if the change in firing rate between laser conditions was significant (Wilcoxon signed-rank, p < 0.05). Green lines / labels on cumulative histograms mark the proportion of units (across all clusters) in which laser presentation suppressed spiking activity.

To capture the distinct effects of light delivery on the neural population, we used K-means clustering to classify units into separate groups based on their responses to sounds with and without laser light. Assessing cluster performance with K between 1 and 10 (see Methods) indicated that two clusters captured the majority of variance between units, with the two groups being distinguished by their sensitivity to photostimulation both when considering auditory evoked activity (**Fig. 4C**) and spontaneous activity (**Fig. 4D**).

When comparing the effects of laser light on sound evoked firing, Cluster 1 showed a significant decrease with photostimulation (n = 272 units, median change of −1.296 spikes/s, Wilcoxon signed rank test with Bonferroni correction, p < 0.001, Z = −14.3), whilst Cluster 2 showed a small but significant increase in firing with light delivery (n = 193 units, median change of 0.0667 spikes/s, p < 0.001, Z = 5.09). In periods of spontaneous activity, Cluster 1 showed a significant decrease in firing with light delivery (median change of −0.4167 spikes/s, Wilcoxon signed rank test with Bonferroni correction, p < 0.001, Z = −8.18), whilst Cluster 2 showed a similar increase in firing in the spontaneous condition as in the evoked condition (median change of 0.0667 spikes/s, p < 0.001, Z = 4.22).

For each unit within a cluster, we also asked if the mean sound-evoked firing rate (windowed between 50 to 150 ms from laser onset, which included 50 ms of baseline activity and the first 50 ms of sound evoked activity) differed between laser presentation and absence (two-tailed sided Wilcoxon signed rank test, p < 0.05). The majority of units in Cluster 1 (60.3 %) showed significant decreases in activity with light delivery, while only a minority of units (25.9%) in Cluster 2 were affected by light delivery. The pattern of results was similar, regardless of whether activity was recorded from single units or multi-units (**Table 2**).

**Table 2:** Proportion of single and multi-units in each cluster that showed a significant change in firing rate with laser light delivery in 50 to 150 ms window after laser onset (Wilcoxon signed rank test, p < 0.05).

|  | Single Unit | Multi-unit | Total |
| --- | --- | --- | --- |
| Cluster 1 | 58 / 103 (56.3%) | 106 / 169 (62.7%) | 164 / 272 (60.3%) |
| Cluster 2 | 16 / 71 (22.5%) | 34 / 122 (27.9%) | 50 / 193 (25.9%) |
| Total | 74 / 174 (42.5%) | 140 / 291 (48.1%) | 214 / 465 (46.0%) |

### Spatial and temporal organization of optogenetic inactivation

The extent and speed of inactivation are major considerations when manipulating neural activity during behavior. To understand how far and how fast it was possible to suppress neurons using ChR2 expressed via the mDlx promoter, we mapped the effects of laser light with cortical depth and time (**Fig. 5**). In our analysis of depth, we defined the limits of auditory cortex on the basis of sound-evoked responses, of which 95% were observed within 2.62 mm of the top of the probe (**Fig. 5A-B**). Such functional estimates are comparable with the thickness of ferret auditory cortex observed histologically (with correction for tissue shrinkage during fixation, **Fig. 3A**).

**Figure 5.**
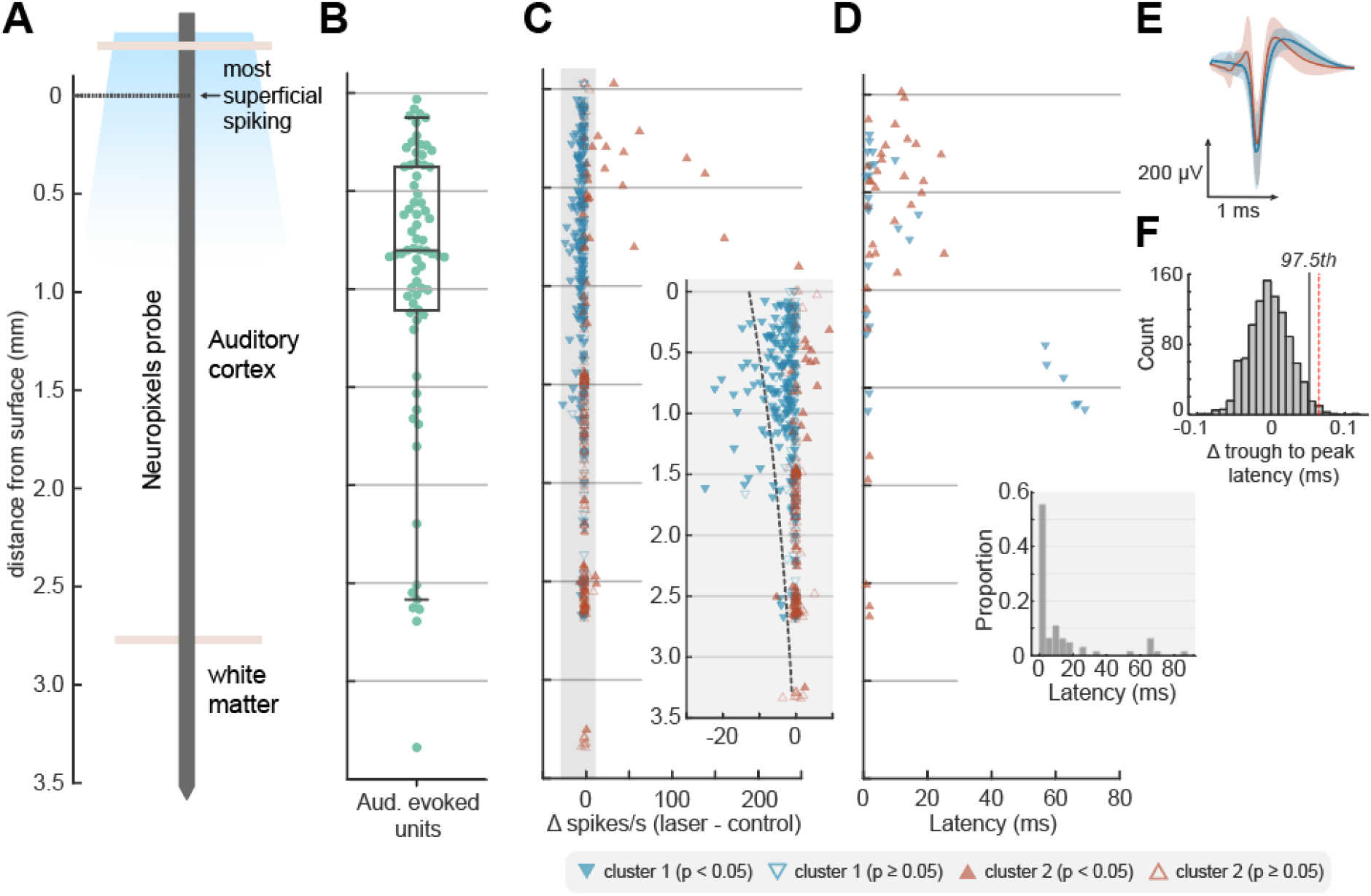
Depth-dependent suppression. (**A**) Schematic of probe displaying approximate anatomical locations in reference to surface calculated by the most superficial depth at which spiking was observed. (**B**) The location of auditory evoked units (n = 72) as a function of cortical depth from surface with boxplot showing quartiles with whiskers showing the 95th percentiles. (**C**) Change in firing rate with light delivery as a function of cortical depth from surface. Inset shows magnified gray region with dotted line showing predictions from fitted Poisson mixed-model. (**D**) Latency of significant change in firing rate with light delivery as a function of depth. Marker color and shape in C-D indicates cluster grouping identified via K-means clustering, as in Figure 4. (**E**) Average spike shapes of well-isolated single-units of cluster 1 (blue, n = 80 SUs) and cluster 2 (red, n = 20 SUs) recorded within 1.598 mm of the surface (error: std.). (**F**) Difference in trough to peak latency of each mean waveform (cluster 1 - cluster 2) for observed data (red dashed line, difference = 0.0648 ms) or when randomly shuffling clusters labels (histogram, n = 1000 iterations) during permutation testing (97.5th percentile, black line).

Across the depth profile of auditory cortex, laser-driven suppression of neural activity was stronger in more superficial units and diminished with distance from the cortical surface (**Fig. 5C**). The effect of depth was evident in the median position of units in clusters 1 and 2 (identified through K-means clustering in the previous section), with light-suppressed units grouped in cluster 1 occurring significantly closer to the cortical surface (rank-sum test, p < 0.001).

Modeling the laser-related change in single trial spike counts of individual units as a function of distance from the cortical surface confirmed a significant interaction between depth and light delivery (Poisson mixed-model regression with distance and light as fixed effects, ferret, unit and sound duration as random effects, p < 0.001). However, the fall-off in suppression captured by the model took place across several millimeters, with 90% of all significantly inactivated units (**Table 2**) being located within 1.598 mm of the cortical surface. This prolonged fall-off over several millimeters contrasts with the rapid attenuation of blue light in tissue over hundreds of micrometers (Li et al., 2019), making it unlikely that light-based artifacts account for the spatial extent of inactivation observed.

The temporal profile of inactivation also indicated that the effects we observed were not a trivial result of cortical heating, as light delivery suppressed cortical activity rapidly (**Fig. 5D**).

Nonparametric cluster statistics revealed a median latency for significant change in firing at 2.5 ms. Such rapid changes in firing rate show that the mDlx-induced expression of ChR2 in auditory cortex provided a fast method for cortical inactivation, and are unlikely to be driven by changes in temperature of tissue that have been reported over longer time-scales, on the order of hundreds of milliseconds or seconds (Owen et al., 2019).

### Optogenetic inactivation primarily affects broad-spiking neurons

Analysis of light-driven suppression of sound driven responses indicated that optogenetic inactivation affected a specific subgroup of neurons; that is units in cluster 1 but not cluster 2, identified through K-means clustering. It is possible that cells within each cluster may be drawn from distinct populations of neurons suppressed by light-driven local network inhibition (cluster 1), and GABAergic interneurons driven by light (cluster 2). Pyramidal neurons and interneurons are often distinguished by their spike waveform as broad and narrow spiking cells respectively (Niell and Stryker, 2008; Moore and Wehr, 2013) and so we asked if the clusters identified from firing rate data might have distinct spike shapes that correspond to these cell types.

To compare spike shapes, we measured the trough-to-peak latencies of average spike waveforms from well-isolated single units in cluster 1 (n = 80) and cluster 2 (n = 20) recorded within 1.598 mm from the cortical surface (i.e. the depth range within which 90% of significantly inactivated units were identified). We found that the trough to peak latencies of single units in cluster 1 (i.e. those that were suppressed by the laser) were indeed longer than those in cluster 2, indicating a broader waveform (**Fig. 5E**).

To determine whether differences in trough-to-peak latency observed between clusters might arise spuriously, we compared the difference we observed in our data with results when randomly shuffling cluster labels (**Fig. 5F**). Permutation testing confirmed that the difference in spike widths between clusters was significant (p = 0.01, n = 1000 iterations). Thus our results are consistent with the suggestion that neurons suppressed by the laser were primarily broad-spiking excitatory/pyramidal neurons, while the remaining cells were more likely to be narrow-spiking inhibitory interneurons. Note however that because the mDlx promotor is specific only to GABAergic neurons, light is likely to drive multiple subclasses of inhibitory interneurons including, but not restricted to, fast spiking PV neurons.

### Auditory cortex is required for vowel discrimination in co-located noise but not clean conditions

We examined the role of primary auditory cortex in behavior using cortical inactivation via cooling in two ferrets (F1311 and F1509) or stimulation of inhibitory interneurons using optogenetics in one ferret (F1706). Ferrets were trained to report the identity of vowel sounds (F1311: /a/ and /i/, F1509, F1706: /u/ and /εe/) of varying sound level in clean conditions (**Fig. 1A**), , and then tested with vowels in additive broadband noise in control conditions and with cooling or laser light delivery.

Auditory cortical inactivation impaired vowel discrimination in noise in each animal (**Fig. 6**). Across SNRs, performance discriminating sounds in noise was worse during cooling than control sessions (change in performance [cooled-control]: F1311 = −11.1%, F1509 = −9.72%) and worse on trials when light was delivered ([Light: On - Off]: F1706 = −12.5%). In contrast, cooling did not impair vowel discrimination in clean conditions in either animal tested (F1311 = +5.39%, F1509 = +1.85%, F1706 not tested with laser light delivery in clean conditions).

**Figure 6.**
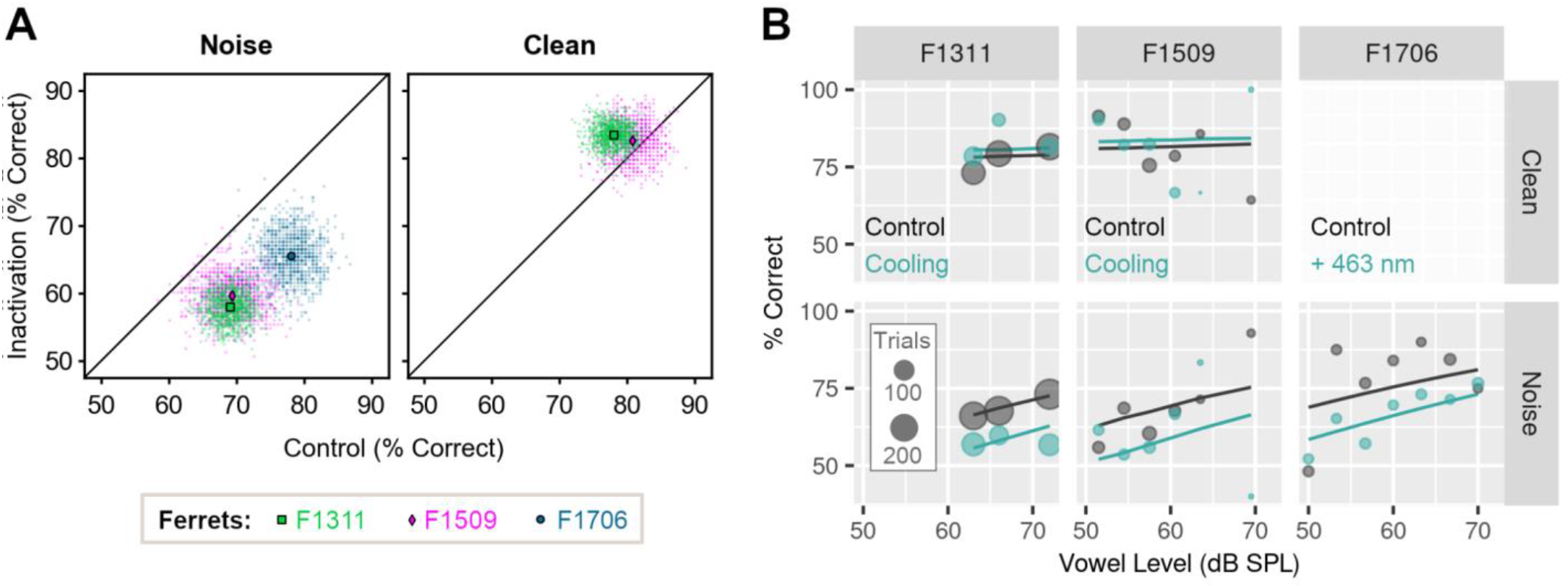
Cortical inactivation impairs vowel discrimination in noise, but not clean conditions. (**A**) Performance discriminating vowels in noise (n = 3 ferrets) or clean conditions (n = 2 ferrets, F1706 not tested) during cooling or optogenetic inactivation and in control testing. Scatter plots show performance across all SNRs or sound levels for each bootstrap (n = 1000 iterations), with means shown as markers. (**B**) Model fit to data (lines) from each ferret discriminating vowels in clean and noise conditions, with cooling (F1311 and F1509) or optogenetics (F1706, noise only). Scatter plots show observed data, with marker size showing trial numbers.

To assess changes in vowel discrimination with cortical inactivation across ferrets, we compared single trial performance using a mixed-effects logistic regression with ferret as a random effect, and in which background noise (clean vs. noise), experimental treatment (test [cooled or light-on] vs. control [warm or light-off]) and the interaction between treatment and noise were contrasted as fixed effects. We also included whether the subject was rewarded at the center spout and the sound level of vowels as covariates, as well as the interaction between sound level and noise condition. Using the Akaike Information Criterion, this model was selected over other alternatives that either omitted interactions, or included three-way interactions between noise, treatment and sound level.

The impairment in vowel discrimination in noise with cortical inactivation was reflected in the fitted model as a significant interaction between noise condition and experimental treatment (**Table 3**, p = 0.002). There was also a significant main effect of noise (p < 0.001) that captured the general impairment of performance caused by degrading sounds. There was no main effect of treatment alone (p = 0.374), illustrating that the general ability to perform a two choice task was not affected by cooling/light delivery.

**Table 3:** Model output for mixed-effect model logistic regression (n = 3 ferrets) showing coefficient estimates and standard error for fixed effects. Sample sizes: F1311 = 1914 trials, F1509 = 603 trials, F1706 = 352 trials. Model fit: Marginal R^2^ = 0.076, Conditional R^2^ = 0.090.

| Fixed Effects | Estimate | Std. Error | Z | P(> z) |
| --- | --- | --- | --- | --- |
| Intercept | 1.473 | 0.266 | 5.538 | < 0.001 |
| Treatment | 0.151 | 0.170 | 0.888 | 0.374 |
| Noise condition | -0.951 | 0.263 | -3.614 | < 0.001 |
| Vowel sound level | 0.112 | 0.325 | 0.344 | 0.730 |
| Center Reward | -0.080 | 0.104 | -0.766 | 0.443 |
| Treatment * Noise | -0.603 | 0.199 | -3.029 | 0.002 |
| Noise * Level | 0.612 | 0.344 | 1.78 | 0.075 |

### Spatial separation of vowel and noise

In the initial vowel discrimination task, vowels and noise were presented together from two speakers; one on the left and right of the head. We also tested a variant of the task in which vowel and noise were presented either together at a single speaker, or spatially separated from left and right speakers (**Fig. 1B**). In initial behavioral testing, we measured the extent of spatial release from energetic masking in six animals: two ferrets implanted with cooling loops (F1311 and F1509) as well as four additional ferrets that were not used for cortical inactivation (F1201, F1203, F1216 and F1217).

Spatial separation of vowel and noise improved the ability of each ferret to discriminate vowel identity compared to colocated vowel and noise (**Fig. 7A**). In terms of percent correct, the benefit of spatial separation was consistent but small for each subject (mean across bootstrap resamples, separated - colocalized; F1311: +1.35%, F1509: +2.85%, F1201: +1.73%, F1203: +2.68%, F1216: 1.94%, F1217 = +2.19%). To relate these results to the maximum unmasking possible, we also measured the effect of removing noise entirely by presenting vowels from a single speaker in clean conditions. Removing noise improved performance (mean across bootstrap resamples, clean - colocalized; F1311: 5.78%, F1509: 15.1%, F1201: 26.6%, F1203: 16.7%, F1216: 11.9%, F1217: 20.2%), but no animal performed perfectly in clean conditions (**Fig. 7B**). Thus, although the absolute changes in performance with spatial separation of noise and vowel were small, they could represent a substantial fraction (up to one fifth) of the behavioral benefit observed when removing noise entirely.

**Figure 7.**
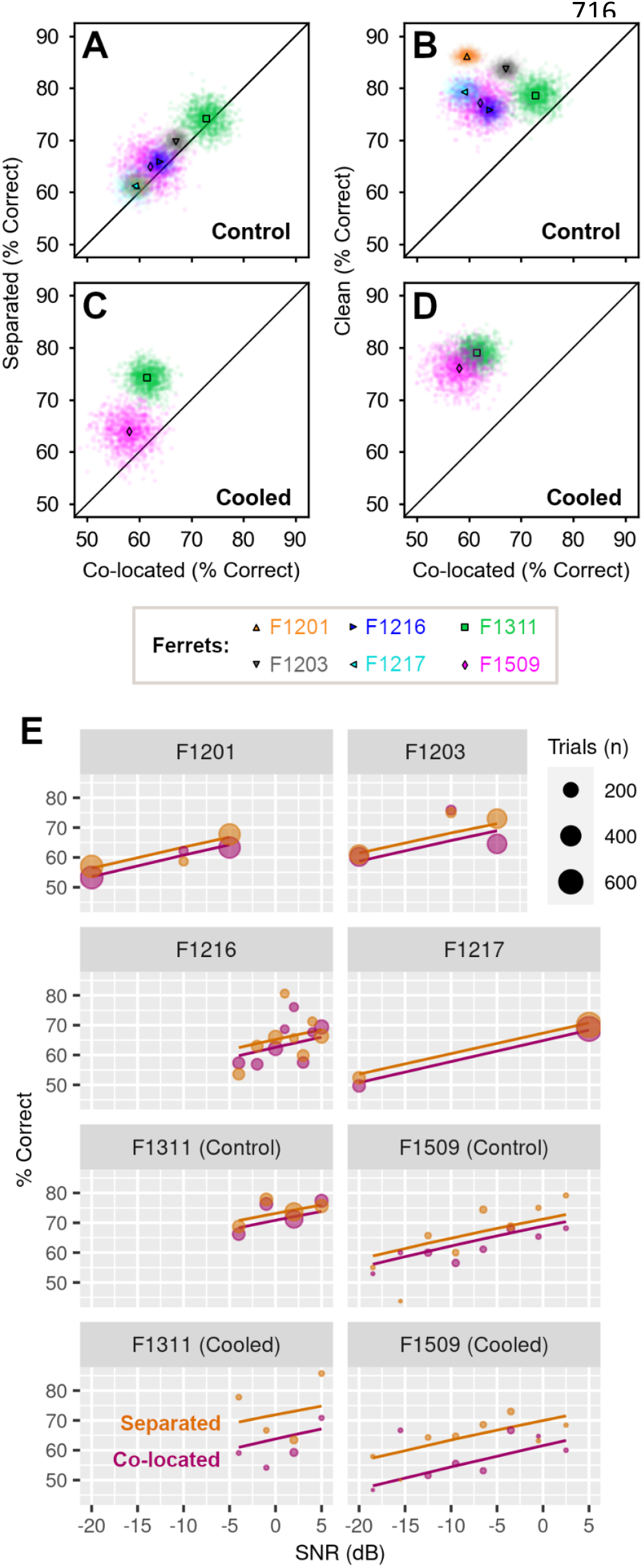
Spatial separation improves vowel discrimination in noise. (**A**) Performance of each ferret (n = 6) in spatially separated or co-located noise in control conditions across SNR. Scatter plots indicate performance across bootstrap resampling (n = 1000 iterations) with mean performance shown by markers. (**B**) Control performance discriminating vowels in clean conditions (i.e. without noise) vs. co-located noise. (**C**) Performance of two ferrets during cooling, when discriminating vowels in spatially separated or co-located noise. (**D**) Performance during cooling when discriminating vowels in clean conditions vs. co-located noise. (**E**) Mixed-effect model fit (lines) and observed performance (markers) vs. SNR discriminating vowels in co-located and spatially separated noise.

Spatial separation also improved vowel discrimination in noise during auditory cortical inactivation. In two animals tested with bilateral cooling, performance was better in spatially separated than co-located noise (**Fig. 7C**, separated - colocalized, F1311: +12.7%, F1509: +5.87%). The benefit of spatial separation was larger during cooling than control conditions (F1311: +1.35%, F1509: +2.85%), primarily because cooling impaired vowel discrimination in co-located noise, and the effect of cooling was ameliorated by spatially separating the vowel and noise. The performance benefit of removing noise completely was also evident during cooling (**Fig. 7D**) and more pronounced than in control conditions ([clean - colocalized], cooled vs. control: F1311: 17.5% vs. 5.78%, F1509: 18.1% vs 15.1%).

To model the effects of spatial separation of vowel and noise on task performance, we fitted a mixed-effects logistic regression to response counts from all animals, with ferret as a random effect and with noise condition (separated vs. co-located), treatment (cooled vs. control), sound level and vowel location (left vs. right) as fixed effects. To account for the possibility that cortical inactivation modulated the effect of spatial separation, we also included an interaction term between treatment and noise condition. Model fitting confirmed the importance of spatial separation (p = 0.009) and the effect of cooling on vowel discrimination in noise (p = 0.011; **Table 4**), as well as the relationship between task performance and sound level (p < 0.001, **Fig. 7E**). There was no significant interaction between cooling and separation, indicating that, at least for the animals tested, cortical inactivation did not affect the performance gained by separating vowel and noise.

**Table 4:** Coefficients for mixed effects logistic regression model comparing vowel discrimination in co-located or spatially separated noise, with cortical cooling (2 ferrets) and in control conditions (6 ferrets). Trial counts: F1201= 3112 trials, F1203 = 2501 trials, F1216 = 2821 trials, F1217 = 2335 trials, F1311 = 2744 trials, F1509 = 1430 trials. Model fit: marginal R^2^ = 0.022, conditional R^2^ = 0.029.

| Fixed Effects | Estimate | Std. Error | Z | P(> z) |
| --- | --- | --- | --- | --- |
| Intercept | 0.173 | 0.087 | 1.98 | 0.047 |
| Noise separation | 0.114 | 0.044 | 2.62 | **0.009** |
| Cooling | -0.322 | 0.126 | -2.55 | **0.011** |
| Vowel sound level | 0.740 | 0.078 | 9.46 | **< 0.001** |
| Vowel Location | -0.031 | 0.042 | -0.747 | 0.455 |
| Cooling * Separation | 0.260 | 0.167 | 1.55 | 0.120 |

### A shared role for auditory cortex in sound localization and vowel discrimination in noise

To determine whether the region of auditory cortex that we inactivated was also involved in spatial hearing, we retrained the two ferrets implanted with cooling loops in an approach-to-target sound localization task (**Fig. 1C**). Sound localization was then tested in control conditions and when cooling auditory cortex bilaterally or unilaterally, with cooling of the left or right auditory cortex only.

Bilateral cooling impaired sound localization in both ferrets (**Fig. 8A-B**), with performance (percent correct) being significantly lower during cooling than in control testing (change in performance [cooled-control]: F1311 = −14.8%, F1509 = −12.9%).

**Figure 8.**
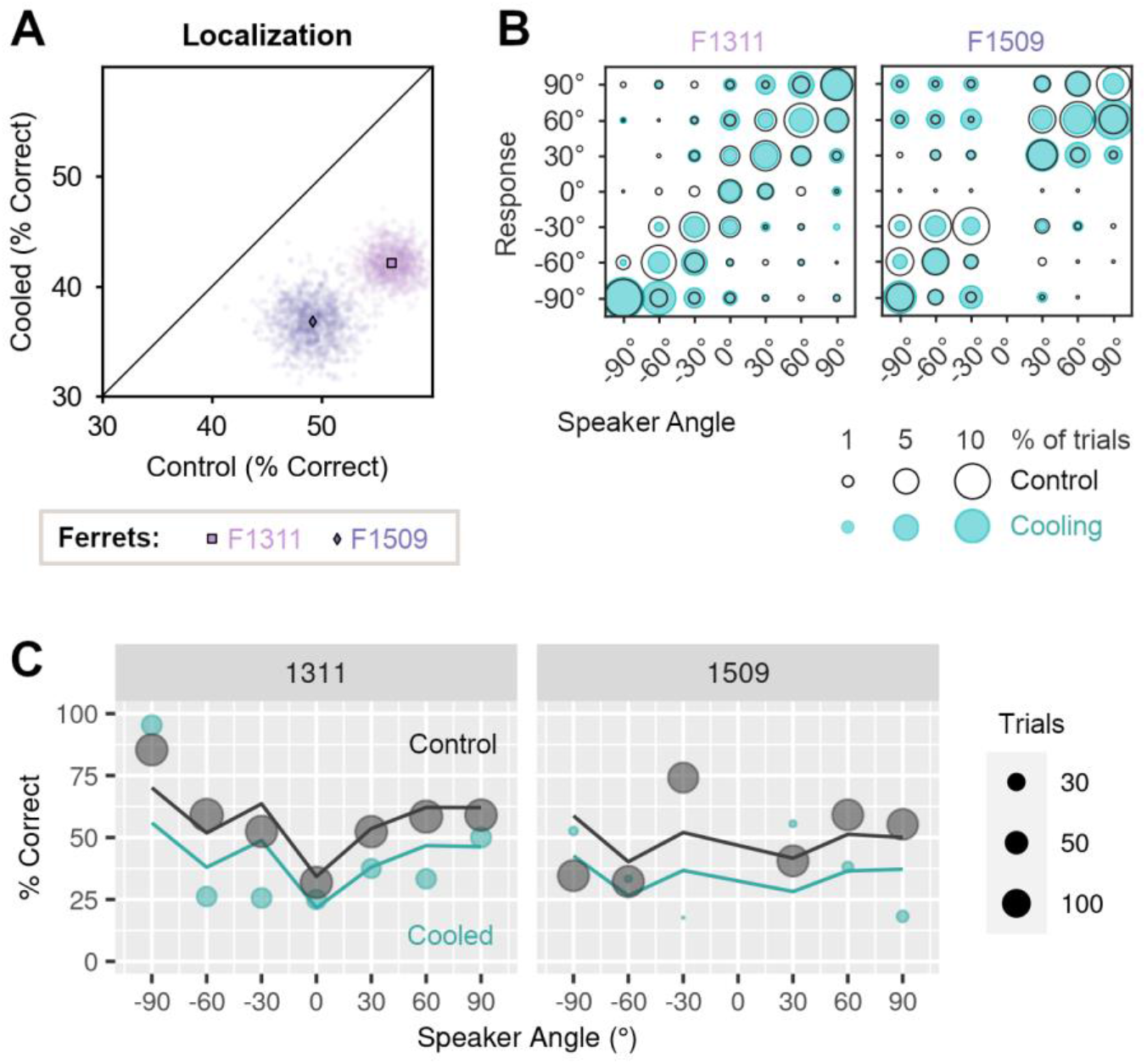
Effects of bilateral cooling on sound localization. (**A**) Performance of ferrets (n=2) tested with bilateral cooling during sound localization. Scatter plots show performance on each bootstrap sample (n = 1000) with means indicated by markers. (**B**) Confusion matrices showing behavioral responses for each speaker and response location in control conditions (unfilled black: F1311 = 1690 trials, F1509 = 1220 trials), and during bilateral cooling (blue: F1311 = 294 trials, F1509 = 115 trials). (**C**) Performance as a function of sound location predicted by mixed-effects logistic regression (lines) and observed during behavior (markers) in cooled and control conditions.

We modeled the effects of bilateral cooling on single trial performance using mixed-effects logistic regression, with treatment [cooled/control] as a fixed effect, and with sound level and center reward as additional covariates in the fixed model. In the random model, we included ferret and speaker location; speaker location was included in the random rather than fixed model to avoid the non-linear dependence of performance on sound location (**Fig. 8C**). The resulting model fit confirmed a significant main effect of cooling, as well as sound level and center reward (p < 0.001, **Table 5**).

**Table 5:**
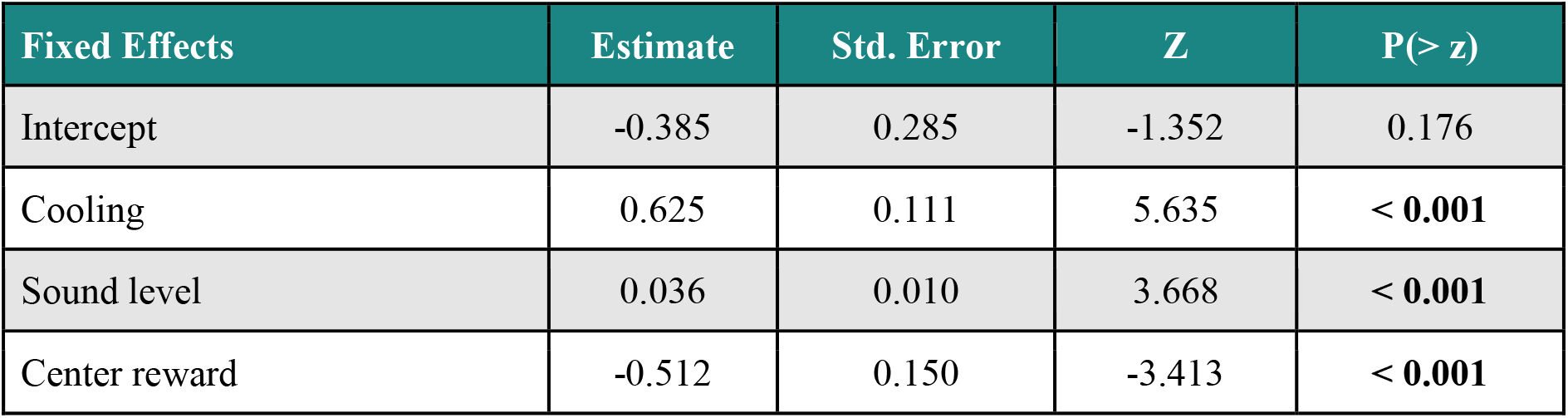
Model results for comparison of performance localizing sounds during cooling and control conditions (n = 2 ferrets). Sample sizes, control conditions: F1311 = 1690 trials, F1509 = 1220 trials, bilateral cooling: F1311 = 294 trials, F1509 = 115 trials. Model fit: marginal R^2^ = 0.039, conditional R^2^ = 0.120.

| Fixed Effects | Estimate | Std. Error | Z | P(> z) |
| --- | --- | --- | --- | --- |
| Intercept | -0.385 | 0.285 | -1.352 | 0.176 |
| Cooling | 0.625 | 0.111 | 5.635 | **< 0.001** |
| Sound level | 0.036 | 0.010 | 3.668 | **< 0.001** |
| Center reward | -0.512 | 0.150 | -3.413 | **< 0.001** |

Unilateral cooling impaired localization of sounds in the contralateral hemifield of space to a greater extent than sounds in the ipsilateral hemifield (**Fig. 9A-B**). Cooling left auditory cortex resulted in larger impairments when localizing sounds in the right side of space, compared to the left (mean change in performance across bootstrap iterations, right vs left speakers, F1311: −19.1 vs + 0.4%, F1509: −32.5 vs −15.6%). Cooling the right auditory cortex had a less detrimental effect, but again resulted in larger deficits in contralateral localization; here, performance was more strongly impaired when localizing sounds in the left than right side of space for one ferret (change in performance, left vs. right speakers, F1509: −22.9 vs. −6.4%). The same pattern of results was observed in the other ferret, but the difference in performance between speaker locations was much smaller (F1311: −8.3 vs −7.5%). In comparison, the effects of bilateral cooling were similar when localizing sounds in both left and right hemifields (F1311: −14.5 vs. −16.6%, F1509: −15.6% vs. −13.4%).

**Figure 9.**
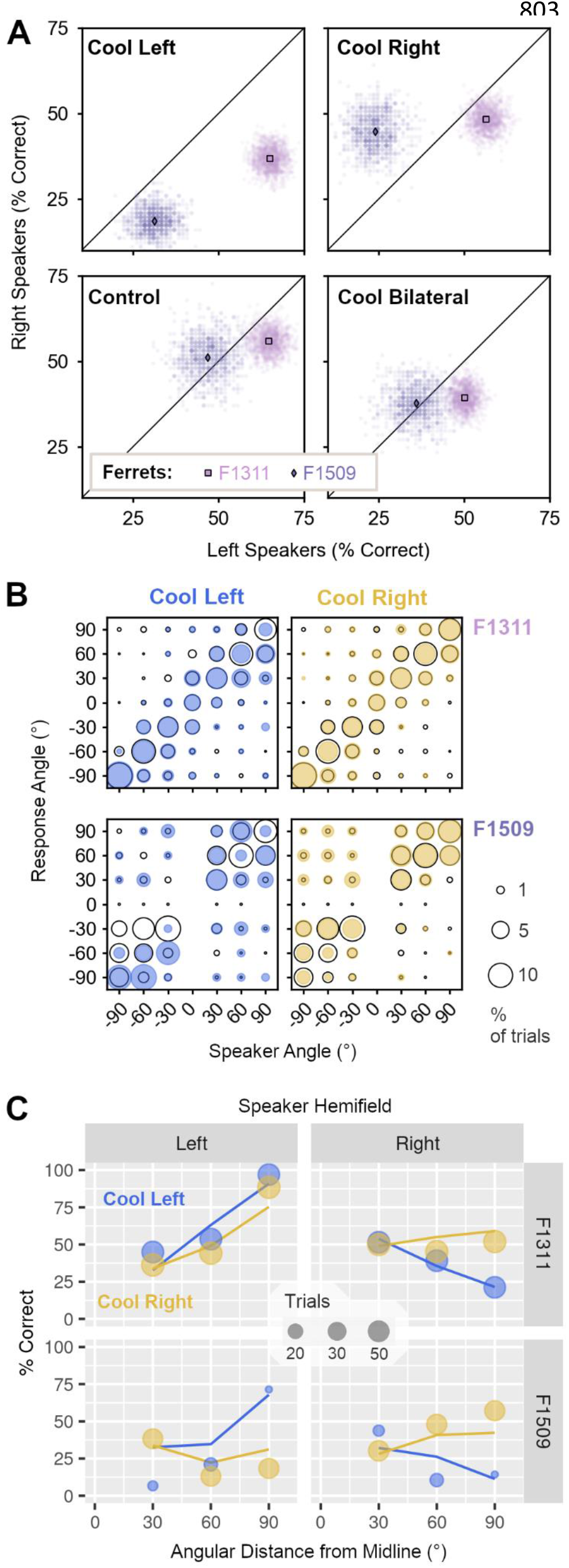
Effects of unilateral cooling on sound localization. (**A**) Performance of ferrets (n=2) localizing sounds in the left and right side of space during unilateral cooling of left or right auditory cortex, control conditions and bilateral cooling. Scatter plots show performance on each bootstrap sample (n = 1000) with means indicated by markers. (**B**) Bubble plots showing the joint distribution of behavioral responses for each speaker and response location during unilateral cooling (filled blue/yellow) and control conditions (unfilled black) for F1311 (top row) and F1509 (bottom row). Sample sizes in control conditions (F1311 = 1690 trials, F1509 = 1220 trials), and during cooling left (F1311 = 476 trials, F1509 = 97 trials) or right auditory cortex (F1311 = 536 trials, F1509 = 294 trials). (**C**) Observed behavior (markers) and model prediction (lines) of performance localizing sounds in left and right sides of space during unilateral cooling.

To model performance during unilateral cooling, we used a mixed-effects logistic regression with ferret as a random effect, and fixed effects for cooled hemisphere (left or right auditory cortex), speaker hemifield (left or right side of space) and distance from each speaker to the midline (30°, 60° or 90°). Comparison of nested models demonstrated that interactions between each parameter, up to the level of the three-way interaction significantly improved model fit (analysis of deviance, p < 0.001) and so we included all interactions between these terms. We also included sound level, the occurrence of a center reward and performance on control trials without cortical cooling (expressed as proportion of trials correct when all other variables were held constant) as covariates.

The model captured the larger effect of unilateral cooling on sound localization in the contralateral hemisphere described above as a significant interaction between cooled hemisphere and speaker hemifield (p = 0.008, **Table 6**). The interaction between cooled hemisphere, speaker hemifield and angular distance of speaker from the midline (p < 0.001) also emphasized how the effects of unilateral cooling were increasingly pronounced at peripheral sound locations (**Fig. 9C**).

**Table 6:** Model results for comparison of performance localizing sounds during unilateral cooling of left and right auditory cortex (n = 2 ferrets). Sample sizes during cooling left (F1311 = 476 trials, F1509 = 97 trials) or right auditory cortex (F1311 = 536 trials, F1509 = 294 trials). Model fit: Marginal R^2^ = 0.185, Conditional R^2^ = 0.208.

| Fixed Effects | Estimate | Std. Error | Z | P(> z) |
| --- | --- | --- | --- | --- |
| Intercept | -3.215 | 0.496 | -6.487 | < 0.001 |
| Cooled Hemisphere | 0.617 | 0.492 | -1.255 | 0.210 |
| Speaker Hemifield | 2.636 | 0.531 | 4.968 | < 0.001 |
| Angle to midline | 2.06 | 0.390 | 5.281 | < 0.001 |
| Sound level | 0.011 | 0.017 | 0.637 | 0.524 |
| Center Reward | -0.082 | 0.296 | -0.276 | 0.782 |
| Control Performance | 2.362 | 0.419 | 5.636 | < 0.001 |
| Hemisphere * Hemifield | -1.769 | 0.669 | -2.642 | 0.008 |
| Hemisphere * Angle | -1.138 | 0.470 | -2.419 | 0.016 |
| Hemifield * Angle | -3.581 | 0.516 | -6.937 | < 0.001 |
| Hemifield * Angle * Hemisphere | 2.965 | 0.632 | 4.694 | < 0.001 |

## Discussion

Our results, summarized in **Table 7**, demonstrate that both vowel discrimination in noise and sound localization depend on a common region of ferret auditory cortex, and that cortical inactivation via cooling leads to behavioral deficits in both tasks, while leaving intact other forms of hearing such as vowel discrimination in clean conditions.

**Table 7:** Summary of results during auditory cortical cooling.

|  | Vowel Discrimination |  |  |  |
| --- | --- | --- | --- | --- |
| Ferret | Clean | CL Noise | SS noise | Sound Localization |
| F1311 | Present | Impaired | Present | Impaired |
| F1509 | Present | Impaired | Present | Impaired |
| F1706 | Not tested | Impaired | Not tested | Not tested |

### Selection of cortical region for inactivation

We implanted cooling loops (or optic fibers) over the MEG, specifically targeting the mid-to-low frequency regions of primary auditory cortex (that border the non-primary fields of posterior ectosylvian gyrus; **Fig. 2A**). We targeted this area as it contains neurons that are predominantly tuned to low sound frequencies (Bizley et al., 2005), often vowel responsive and/or spatially tuned (Bizley et al., 2009; Town et al., 2017, 2018) and may play an important role in encoding interaural timing cues supporting sound localization (Wood et al., 2019). It is thus perhaps unsurprising that a region implicated in processing of spatial and non-spatial sound features should contribute to multiple forms of hearing.

The extent of inactivation is a critical consideration in any perturbation study (Slonina et al., 2022); the size of cooling loops used here reflected a compromise between the need to inactivate sufficient numbers of neurons to observe behavioral deficits, and avoid unintended spread of cooling to subcortical structures (Coomber et al., 2011). Previous data from our lab has shown that the cooling loops we used induce spatially-restricted heat loss that limits the reduction in spiking activity to the cortical layers surrounding the loop (Wood et al., 2017). In the current study, ferrets could discriminate vowels in clean conditions during bilateral cooling, while in the same sessions, vowel discrimination in noise was impaired. The ability of animals to discriminate vowels in clean conditions demonstrates that the cooling protocol we used did not affect ferrets’ general hearing, motor ability or capacity to engage in behavioral tasks.

A critical outstanding question is to what extent non-primary regions of auditory cortex beyond MEG contribute to sound localization and vowel discrimination in noise. Earlier cooling studies have used multiple loops to identify distinct contributions of non-primary areas of cat auditory cortex to spatial and non-spatial hearing (Lomber and Malhotra, 2008). If such distinctions also exist in ferrets, then one would predict that inactivation of distinct fields of non-primary auditory cortex may disrupt specific tasks. Testing this will be an important issue for future investigations, which will benefit from the optogenetic techniques that we have confirmed here are effective in rapidly suppressing auditory cortical processing of sounds and disrupting psychoacoustic task performance.

### What is auditory cortex doing?

Our results confirm the widely observed role of auditory cortex in sound localization in carnivores (Kavanagh and Kelly, 1987; Smith et al., 2004; Malhotra et al., 2008), while the ability of ferrets to discriminate vowels in clean conditions is consistent with similar behavior in cats with lesions of primary and secondary auditory cortex (Dewson, 1964). Thus, although auditory cortical neurons are strongly modulated by vowel timbre (Bizley et al., 2009), there may be redundant encoding of spectral timbre across multiple cortical fields, or this activity may not be required for the simple two-choice timbre discrimination employed here.

A role for auditory cortex in vowel discrimination became evident when we added noise to vowels. An open question from our work is whether the same role for auditory cortex would be observed in clean conditions if vowels were presented closer to ferret’s psychophysical thresholds. If so, then our current results might indicate a role for auditory cortex in difficult listening conditions that is consistent with deficits in fine spectrotemporal discriminations following auditory cortical lesions in cats and non-human primates that otherwise have limited effects on easier tasks requiring coarser resolution (Evarts, 1952; Goldberg and Neff, 1961; Diamond et al., 1962; Massopust et al., 1965; Dewson et al., 1969; Heffner and Heffner, 1986). Interpreting lesion studies requires caution, due to the potential for recovery of function; however our results were obtained using reversible methods for which there was minimal opportunity for recovery during cooling, and particularly during rapid optogenetic inactivation.

The benefit of spatial separation of vowel and noise to vowel discrimination observed during cooling, coupled with impaired sound localization is consistent with separate mechanisms for spatial stream segregation and discrimination of sound location (Middlebrooks and Onsan, 2012). That spatial separation of target vowels and noise maskers benefits vowel discrimination during cortical cooling suggests that subcortical structures can parse the noise and vowel into separate streams. In stark contrast, substantial performance deficits were observed for co-located vowel and noise, emphasizing the importance of auditory cortex in segregating competing co-located sound sources (Mesgarani and Chang, 2012; Bizley and Cohen, 2013). In our results, the spatial separation of target and masker into opposing hemifields may result in the representation of the vowel being comparable to that in clean conditions in the hemisphere contralateral to the vowel, and help animals to compensate for the lack of cortical scene analysis that is critical for resolving co-located sound sources.

### Spatial release from energetic masking

The effects of release from energetic masking that we observed were small, relative to the benefit of removing masking entirely. This is not surprising given the limited effectiveness of spatial release from energetic masking relative to release from informational masking that have been widely reported (Brungart, 2001; Jones and Litovsky, 2011). It is likely that the benefit animals received from spatial separation of vowel and masker can be accounted for by the better-ear effect, in which spatial separation elevates the signal-to-noise ratio at one ear (while decreasing the SNR at the opposite ear), and listeners are then able to select information available from the better ear. Such effects may arise by the level of the inferior colliculus (IC)(Lane and Delgutte, 2005), which would, at least in part, explain how ferrets retained a benefit of spatial separation during cortical cooling. One would therefore predict that IC inactivation might result in more effective disruption, particularly when inactivating IC contralateral to the better ear. While cooling such deep-lying structures within the ferret brain would likely affect surrounding brain regions, the potential anatomical specificity of optogenetics makes such experiments feasible in the future.

### Auditory decision making

A notable feature of our results, along with the general pattern in the literature on hearing impairments following auditory cortical inactivation, is the preserved ability of animals to perform some sound-based tasks (e.g. vowel discrimination in clean conditions). These findings suggest that substantial redundancy in the auditory system allows alternative pathways to support task performance. The most obvious candidates for this are the ascending pathways from medial geniculate thalamus to secondary auditory cortex that bypass primary fields of the MEG (Bizley et al., 2015).

It is also possible that information at earlier stages of the auditory system, in this case about vowel identity (Blackburn and Sachs, 1990; Schebesch et al., 2010), can access brain areas that coordinate behavior and is sufficient for discriminations that have already been learnt (Ponvert and Jaramillo, 2019). Our use of reversible inactivation via cooling, which operates on the timescale of minutes / individual test sessions, suggests that any redundant pathways must come into use rapidly, integrate seamlessly with normal decision-making processes and occur with minimal need for learning.

Understanding how signals in auditory cortex are integrated into behavior is also critical for determining how deficits in spatial and non-spatial hearing arise, as the impairments observed in vowel discrimination in noise and sound localization may not have arisen through the same mechanisms. Cooling suppresses the activity of neurons, and so we might infer that the absence of spiking degrades cell assemblies that downstream neurons rely on for informed auditory decision making. Such downstream centers may be located in areas such as the prefrontal cortex (Romanski et al., 1999; Kaas and Hackett, 2000) or the striatum (Znamenskiy and Zador, 2013). To ascertain the underlying causes of the deficits we have observed, it will be necessary to combine auditory cortical inactivation with neural recording in such downstream areas, or to perform targeted manipulations of specific neural pathways.

### A role for areas showing mixed selectivity in perception?

We targeted inactivation to the area of auditory cortex in which neurons have previously shown mixed selectivity for sound location and vowel identity (Bizley et al., 2009; Walker et al., 2011; Town et al., 2018). Such mixed selectivity has been observed widely, including across the auditory system (Cohen et al., 2004; O’Connor et al., 2010; Chambers et al., 2014; Downer et al., 2017; Yi et al., 2019; Amaro et al., 2021) and may reflect a general process through which neural systems meet the demands of complex and flexible behaviors (Rigotti et al., 2013; Jazayeri and Afraz, 2017). Our results show that an area of the brain tuned to multiple sound features makes a contribution to multiple forms of hearing, and are consistent with broader predictions about the involvement of mixed selectivity in behavior (Fusi et al., 2016).

Mixed selectivity expands the range of dimensions across which groups of neurons can represent sounds, and so it may be possible to recover detailed information about diverse stimulus sets from population activity in auditory cortex. However, our ability to observe the use of such information in animal behavior is still limited, as most behavioral tasks are low-dimensional (i.e. they have only one or two independent variables along which subjects act)(Gao and Ganguli, 2015). By testing the effects of cortical inactivation on both spatial and non-spatial hearing in the same subjects, we have taken some of the first steps towards expanding the study of auditory behavior to higher dimensions that may be necessary to understand the role of mixed selectivity in everyday hearing.

## Data Availability

All code and data associated with the project is available at: https://github.com/stephentown42/cooling_auditory_cortex

## Acknowledgements

We would like to thank Dr Tara Etherington for assistance with data collection during cortical cooling and Dr Joseph Sollini for assistance in developing the optogenetic approach. We are also grateful to Dr Erwin Alles for constructing the fiber optic implants used with F1801 and F1807 and Linda Ford for logistical support.

Data associated with sound localization of one subject (F1311) has been previously reported in (Wood et al., 2017)

This research was funded in whole, or in part, by the Wellcome Trust [a Wellcome Trust / Royal Society Sir Henry Dale Fellowship to JKB, Grant number 098418/Z/12/A], a Royal Society Dorothy Hodgkin Fellowship to JKB, the BBSRC (BB/H016813/1) and the European Research Council (SOUNDSCENE). For the purpose of open access, the author has applied a CC BY public copyright license to any Author Accepted Manuscript version arising from this submission.

